# Catalyzing computational biology research at an academic institute through an interest network

**DOI:** 10.1101/2025.01.05.631403

**Authors:** Jaroslav Zak, Ian Newman, Daniel J. Montiel Garcia, Daniele Parisi, Janet Joy, Steven R. Head, Jean-Christophe Ducom, Padma Natarajan, Haissi Cui, Sabah Ul-Hasan

## Abstract

Biology has been transformed by the rapid development of computing and the concurrent rise of data-rich approaches such as -omics or high-resolution imaging. However, there is a persistent computational skills gap in the biomedical research workforce. Inherent limitations of classroom teaching and institutional core support highlight the need for accessible ways for researchers to explore developments in computational biology. An analysis of the Scripps Research Genomics Core revealed an increasingly diverse set of experiments: the share of experiments other than bulk RNA- or DNA-seq increased from 34% to 60% within 10 years, requiring more tailored computational analyses. These challenges were tackled by forming a volunteer-led affinity group of over 300 academic biomedical researchers interested in computational biology referred to as the Computational Biology and Bioinformatics (CBB) affinity group. This adaptive group has provided continuing education and networking opportunities through seminars, workshops and coding sessions while evolving along with the needs of its members. A survey of CBB’s impact confirmed the group’s events increased the members’ exposure to computational biology educational and research events (79% respondents) and networking opportunities (61% respondents). Thus, volunteer-led affinity groups may be a viable complement to traditional institutional resources for enhancing the application of computing in biomedical research.

## Introduction

Since the mid 1960s^1^ biology has gradually transformed into an information science. Over this period, the volume of biomedical literature has increased exponentially and has been accompanied by an explosion of - omics data. Both the volume of scientific publication^2^ and -omics data^3^ are increasing exponentially. For example, public databases of major types of genomic and proteomic data, such as dbSNP, EGA and ENA have experienced double-digit percent annual growth in the number of records and sequences^4^. Similarly, the number of tools published to analyze -omics data has experienced faster than linear growth^5^. Deep learning based approaches have resulted in paradigm shifts in the area of protein prediction, raising expectations for their transformative effects in other areas of biology, such as drug discovery^6^.

As interaction with large biomedical data has become ubiquitous across research disciplines, so too has the rate of interaction between computational tools and researchers who would otherwise not describe themselves as computational biologists^7^. Without consistent efforts to stay informed, they risk using outdated methods or misapplying tools, resulting in inefficiencies and errors. For instance, the authors of TopHat have noted that TopHat continues to be used for RNA-seq read alignment more each year, nearly ten years after its publication, while superior versions have long been available (first TopHat2, then HISAT, then HISAT2)^8^. Similarly, default settings in software such as Microsoft Excel have led to preventable errors, such as converting gene names into calendar dates, causing data loss ^9, 10^. These issues underscore the need for both awareness of new tools and appropriate training to use them effectively.

Traditionally in academic settings a large part of the dissemination and training for new techniques happens amongst colleagues within the same laboratory group or department. This mechanism can be less effective in multidisciplinary groups^11^, where computational tools may not be the central focus. Institutional strategies for alleviating this challenge have included outsourcing computational analyses to computational biology cores, encouraging collaboration outside of the department (e.g., with computational biology laboratory groups), and offering formal training courses. Yet, significant disparities in computational literacy and utilization persist.

There is a long-standing gap between the expected and actual levels of computational literacy in the biomedical research workforce. Tan et al. proposed a minimum set of skills for university graduates to meet the informatics needs of the ‘-omics era’, including the command of bioinformatic tools and a basic understanding of programming languages^12^. A survey of 1260 faculty by the NSF Network for Integrating Bioinformatics into Life Sciences Education (NIBLSE) defined core competencies for undergraduate students in the life sciences which include accessing genomic data, genomic tools, some experience with command line tools and writing simple scripts^13^. However, these core competencies are not universally held by researchers as the divide between ‘dry lab’ and ‘wet lab’ biologists persists. A recurring recommendation is for the education curricula to be updated to match the demands of contemporary biology^14^, but formal education alone may not be able to keep pace with rapidly evolving computational technologies nor can it be expected to deliver the tailored information needed for individual research projects. Further, as technological advances continue beyond the completion of formal undergraduate and graduate classes, there is a need for efficient ways of updating computational knowledge and skills for graduated practitioners. Although empirical data on the skills gap in the biomedical research job market is lacking, data from other fields suggest evolving methods create demand for latest computational skills such as data science^15, 16^.

The adoption of computational tools in biomedical research often mirrors broader trends in technology adoption, leading to near-universal uptake in some areas. For example, the National Institutes of Health no longer accept competing grant applications on paper^17^. Electronic slides prepared in presentation software have become the universal format of scientific conferences, and electronic classroom tools have also reached near universal adoption^18^. Similar trends are observed in other sectors, for example, electronic health records are gradually replacing paper copies nationwide^19^. Recent advances in artificial intelligence (AI), including interactive chatbots such as ChatGPT, suggest that AI-powered tools could be headed towards near-universal adoption in the long term. Indeed, in the 3 months following the public release of ChatGPT, an estimated 10% of scientific papers were completed with the assistance of ChatGPT or similar LLM tool, a striking statistic given that many papers published during this time were peer reviewed before ChatGPT was publicly released^20^. This suggests that adoption is often a question of timing rather than inevitability. Together, the skills gap and the rapid emergence of new technologies underscore the need for accessible ways to explore the intersection of computational biology with other disciplines, fostering opportunities to apply computational methods in biomedical research through skill development or collaboration. This broader need for ‘continuing computational education’ goes beyond mastering specific skills and includes maintaining an awareness of the developments in computational biology and opportunities for its application to other disciplines.

Other fields, such as medicine, offer models for structured continuing education through mechanisms like mandatory Continuing Medical Education (CME) requirements. In contrast, continuing academic education requirements are typically optional, leading to wide variability in computational expertise. There is currently no consensus on the most effective mechanisms of continuing education in computational biology for the biomedical research workforce. At the Scripps Research Institute, an independent academic biomedical research institute that also supports a graduate program, we created the Computational Biology and Bioinformatics (CBB) affinity group, a trainee-led community for the discussion of computational biology/bioinformatics with the goal of facilitating knowledge exchange. This study examines how the CBB group at Scripps Research serves as a model for addressing the above challenges, exploring CBB’s role in fostering computational literacy, promoting tool adoption, and bridging training gaps in a multidisciplinary setting.

## Results

### Assessing computational biology activity at an academic biomedical research institute

Despite the pervasive implementation of computational methods in the life sciences, quantifying the degree of computational research activity remains challenging. To gain a better understanding of computational research activity at Scripps Research, we examined official and empirical data from independent sources. As of January 2025, the institute’s website lists 24 (16%) of its 153 faculty as performing research that ‘contains Computational Biology or Computational Biology/Bioinformatics’ and 30 faculty members at the rank of Assistant Professor or above are affiliated with the Integrative Department of Structural and Computational Biology. However, the majority of these faculty also perform research in other areas. Additionally, many laboratories at Scripps rely heavily on computational methods, utilizing advanced analyses in proteomics, genomics, neuroscience, structural biology, virology etc. without being classified as having a focus on Computational Biology. Therefore, we sought a more empirical way of assessing computational research activity at the institute.

We identified next-generation-sequencing-assisted technologies as a major area requiring computational analysis. In addition to sequencing services, the Scripps Research Genomics Core offers services including genomic and transcriptomic library preparation. We examined usage of the Genomics Core over a 10 recent year period using database records. In a dataset spanning the years 2012-2021, an increase in the number of unique experiments performed in the Core was observed: the annual number almost tripled between 2012 and 2018 (**Fig. 1A**).

**Figure 1.**
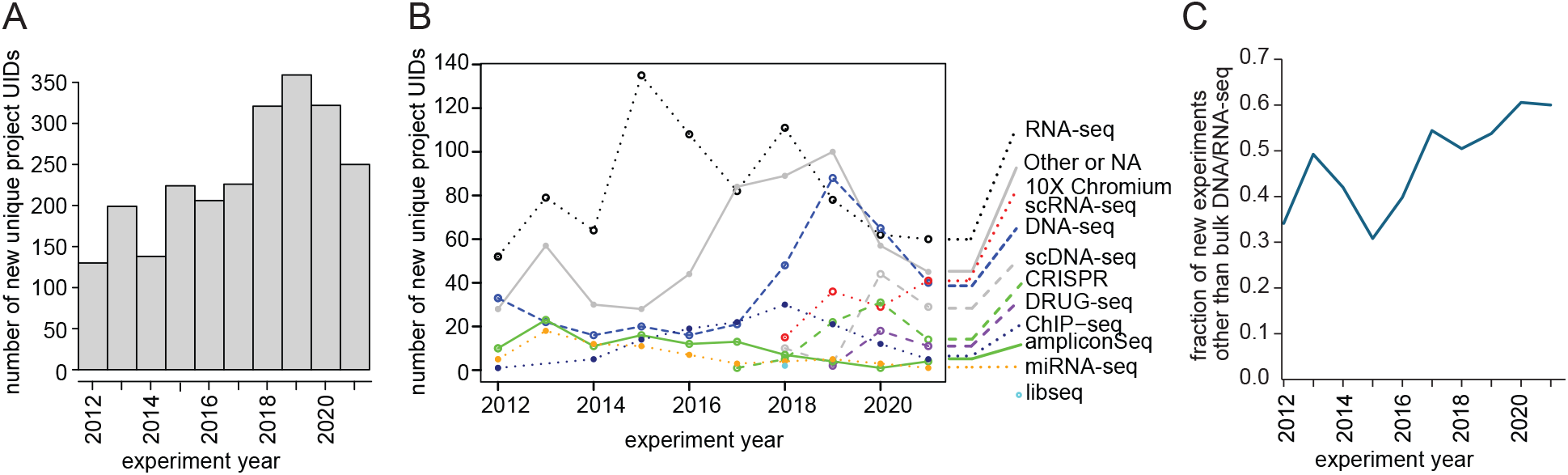
Trends in the usage of the Genomics Core. (A) Overall increase in unique experiments performed in the Genomics Core between 2011-2021. (B-C) Diversification in experiment types performed in the Genomics Core between 2012-2021.

Analyzing the types of experiments performed in the core, bulk RNA-sequencing (RNA-seq) dominated as the most frequent library preparation type in most years (**Fig. 1B, Suppl. Table 1**). However, there was a clear change in the overall composition of experiment types over time: the relative share of bulk RNA-seq decreased from a peak of 60% in 2015 to 24% in 2021 and new techniques such as 10X Chromium single-cell RNA-sequencing (scRNA-seq), CRISPR experiments and high throughput RNA-seq (such as Digital RNA with pertUrbation of Genes (DRUG-seq)^21^) were introduced (**Fig. 1B, Suppl. Table 1**). The drop in the bulk RNA-seq share is only partially explained by the rise in scRNA-seq to 16.4% in 2021; the total diversity of experiment types increased significantly. Accordingly, the fraction of new experiments other than bulk RNA/DNA-seq rose from 34% in 2012 to 60% in 2021 (**Fig. 1C**). Overall, these data suggest that the number of unique experiments as well as the diversity of experiment types in the Genomics Core increased from 2012-2021, reflecting an institutional growth of activity in this research area.

We next examined aggregated usage statistics of the Scripps Research High Performance Computing Cluster (HPC), a shared resource freely available to all institute laboratories on request. There was a slight increase in the total number of unique laboratories utilizing the HPC from 2017 to 2020 (**Table 1**). Notably, the these numbers suggest that the majority of Scripps ∼150 laboratories utilized HPC at least once during the years examined (**Table 1**). We observed no significant increase in the number of central processing unit (CPU) hours per year; however, given improvements in software and hardware performance, this parameter may not appropriately capture overall computing activity. Interestingly, laboratories utilizing the HPC belong to all Scripps Research departments including Chemistry, The Scripps Research Translational Institute and Molecular Medicine (**Table 2**). Molecular Medicine was the top user by CPU hours during the period examined, highlighting that computational biology is not constrained to computational departments. This is at least partly driven by the CPU-heavy algorithms used for proteomics analysis. Multiple departments were also represented among labs whose usage increased the most year-to-year (**Suppl. Tables 2-3**). Importantly, some research groups at Scripps utilize private computational resources such as group-only clusters or cloud computing providers and usage data on those resources were not available for analysis. Therefore, trends in total computational usage might differ from those of HPC usage. Overall, these data reveal an active user base of genomics and computing resources at Scripps Research, spanning multiple departments.

**Table 1.** Quantification of unique research groups using HPC at Scripps between 2017-2020. HPC, high-performance computing cluster.

| Year | Number of unique labs using HPC:<br>Number (approx. % of total laboratories) |
| --- | --- |
| 2017 | 83 (54%) |
| 2018 | 82 (54%) |
| 2019 | 85 (56%) |
| 2020 | 92 (60%) |

**Table 2.**
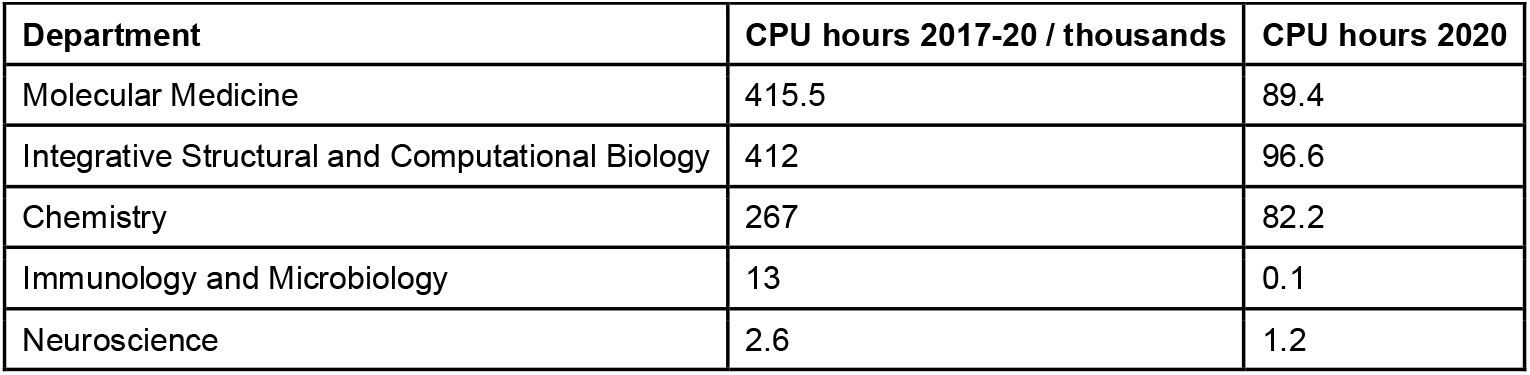
Top departmental users of Scripps HPC 2017-20. NB: These totals include only labs for which data from both 2017 and 2020 were available. This is to remove the effect of labs moving which can distort data. CPU, central processing unit; HPC, high-performance computing cluster.

### Supporting computational research and education at a biomedical research institute

Scripps Research offers data analysis services through its Center for Computational Biology and Bioinformatics (CCBB), a core facility. While the analysis services offered are diverse (**Suppl. Table 4**), the ones best utilized are predominantly genomic analyses. Although a database of specific CCBB service usage is not maintained, the approximate usage of CCBB services between April 2022 and Jan 2023 was reported to comprise mainly quality control for RNA-seq data, subsequent analysis of these datasets, and analysis of ChIP-seq datasets. Noted was also an increase in the number of scRNA-seq analyses conducted. Thus, basic analyses for the most common data types generated in the Genomics Core are supported through the CCBB core whereas niche and iterative analyses are typically performed by individual researchers.

The Skaggs Graduate School of Chemical and Biological Sciences at Scripps Research has consistently offered courses focused on teaching basic computational and bioinformatics skills for the past decade (**Fig. 2, Suppl. Table 5**). Courses range from basic introductions to coding and statistical testing in R to more advanced topics of genetics and genomics and application of advanced statistical models. However, the number and content of these courses are limited to certain areas of computational biology such as statistics (**Fig. 2, Suppl. Table 5**). To supplement these courses, the graduate studies office has supported students pursuing external courses, yet the number of students taking advantage of this benefit is very low (fewer than 5 per year).

**Figure 2.**
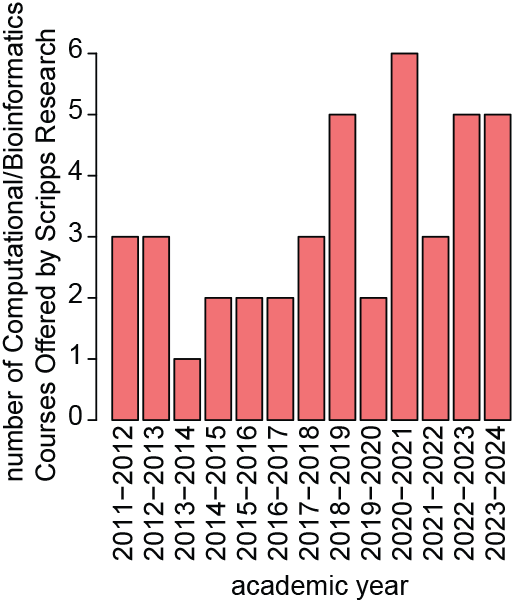
Scripps Research Skaggs Graduate School of Chemical and Biological Sciences course offerings covering computational/bioinformatic topics per year between 2011-2024. Total number of unique courses offered per year. Full list given in Supplementary Table 5.

Thus, institutional support for common data generation, analysis and computational skills training is available. However, given the breadth of departments engaged in computational research and the increasing diversity of experiments performed at the Genomics Core, many scientists still must rely on asking colleagues and self-teaching to analyze their data.

### Enhancing computational research through an interest network

Our data show the scale of computational work at Scripps exceeds the formal learning opportunities present. To address this gap as well as provide a platform for information exchange, collaboration and inter-departmental networking, we initiated the Computational Biology and Bioinformatics (CBB) interest network. The network is run by early-career scientist volunteers and aims to disseminate knowledge related to computational biology, lower the barriers to computational research and connect scientists across disciplines and departments. To achieve these goals, the CBB has performed the following actions.

First, an active directory of members interested in computational biology at all levels of seniority has been compiled to ease collaboration and maintain a census of the computational community at Scripps Research and guests from neighboring institutions. Second, a regular seminar series was launched featuring a broad range of topics connected by some degree of utilization or development of computational methods. To provide opportunities for early-career scientists, speakers from all levels of seniority have been invited and self-nominations were welcomed. To date, 34 seminars featuring speakers from over 10 institutions have been held (**Table 3**). Third, seasonal workshops led by volunteers have been organized to introduce participants to specific topics of interest such as SQL, basics of Python grammar, etc. Fourth, coding sessions featuring code analysis, live presentation, troubleshooting, software package discussion, etc. were held intermittently.

**Table 3.** Overview of CBB activities. Official events organized under the CBB umbrella. GNU, Gnu’s Not Unix; SQL, structured query language.

| Activity | Number held since Feb 2020 | Topics covered | Goals |
| --- | --- | --- | --- |
| Seminars | 34 | Summary in Figure 3. | Disseminate knowledge in computational biology.<br>Share research updates.<br>Facilitate collaborations.<br>Offer presentation opportunities for early-career researchers. |
| Coding sessions | 13 | Object-oriented programming in R, lm() and glm() modeling, CBB Github repo, Developing a Python package, Basic GNU parallel and python multiprocessing, Automating the CBB Github repo, Standardization of visualizing scientifically accurate RNA-seq, CodeWave: AI/ML working group; Deep learning and its applications to research | Exchange programming expertise.<br>Keep up to date with the latest software packages, statistical methods relevant to biomedical research. |
| Workshops | 2 | General code architecture, SQL (3 classes) | Provide introductory training not covered by formal courses at Scripps. |

The seminar series has operated continuously since its initiation in 2019. We examined the seminar topics to understand the range of fields represented in the series. Analyzing the top 3 keywords representing topics of interest per seminar, the most frequent keywords were genomics, machine learning and immunology (**Fig. 3**). However, structural biology, neuroscience and translational science (“therapeutics”) were well represented, indicating that the seminar series transcends traditional disciplinary boundaries (**Fig. 3**). The speakers represent 10 institutions, 6 departments at Scripps Research and span multiple levels of seniority from graduate student to faculty (**Suppl. Table 6**.).

**Figure 3.**
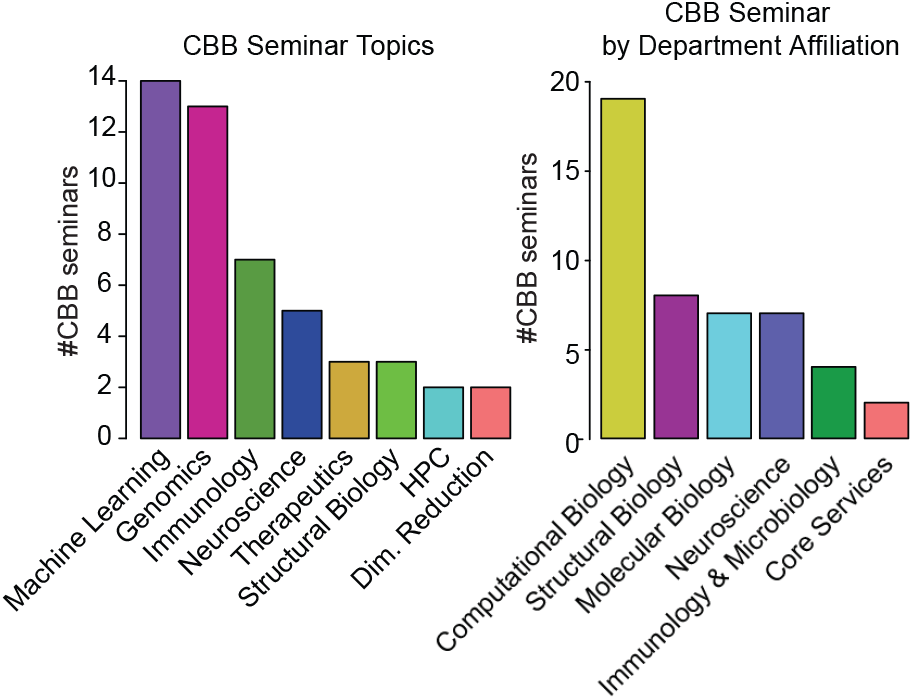
Topics and primary disciplines of CBB seminars. For speakers with unknown department affiliation, department affiliation of the lab principal investigator was used. Department names were grouped by categories of research, e.g., neurology and neuroscience grouped under Neuroscience. Scientific topic for non-Scripps speakers with unknown department affiliation was designated by their lab website landing page as best estimate of the broad research category. Only keywords with at least 2 independent seminars were included in the frequency analysis. There were 5 singleton keywords and 1-3 keywords were used per seminar in the seminar topics analysis.

The seminars were initially held in person but have continued in a virtual format since 2020. The median attendance as of the latest data cutoff (Nov 2023) is 24, with a trend towards increased attendance between 2021 and Nov 2023 (**Fig. 4**). Therefore, the CBB seminar series has attracted consistent participation in its 3.5-year operation.

**Figure 4.**
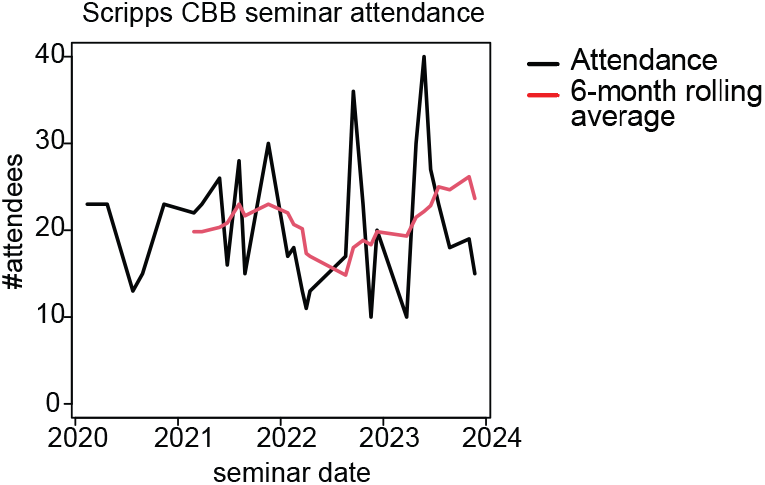
CBB seminar attendance. Attendance at virtual seminars was measured as the number of unique users joining each virtual meeting.

To advance the goal of connecting scientists interested in computational biology and exchanging knowledge and skills in this area, the CBB network organized additional activities (**Table 3**). A member directory and a Slack workspace were compiled to enable efficient contact between interactive coding sessions (held in person or virtually) and to introduce attendees to fundamental programming principles and tools directly applicable to biomedical research. Topics among others included object-oriented programming in R, linear models, how to develop a Python package and advances in large language models. These coding sessions were led by CBB member volunteers, were interactive, featured opportunities for community code troubleshooting and generally did not overlap with TSRI Graduate School courses or other formal education events on campus. This format allowed attendees to interactively learn about topics beyond what is available on campus and seek feedback on potential real-life research applications. Notably, the attendance was particularly strong during the pandemic year of 2021, highlighting the potential of interactive remote coding sessions as a means to maintain productivity when in-person research opportunities are restricted.

### Measuring the impact of the interest network

To assess the impact of Computational Biology and Bioinformatics (CBB) events on the Scripps Research community, we conducted a survey among participants in CBB initiatives. The survey was conducted over 2 months, administered electronically via email invitations to CBB members and past speakers and collected in a de-identified manner. Participants were informed the results may be used for research purposes. All CBB members (approximately 300) received the survey invitation. A total of 28 individuals participated in the survey, an estimated participation rate of 9%.

The first question in our survey, “How many CBB-organized events have you attended?” revealed a substantial engagement. The majority of respondents reported attending 1-3 events (13 participants), followed by 4-6 events (7 participants), and 7 or more events (4 participants). Bias in survey participation is reflected in that only two respondents indicated that they had never attended any CBB events (see **Fig. 5**).

**Figure 5.**
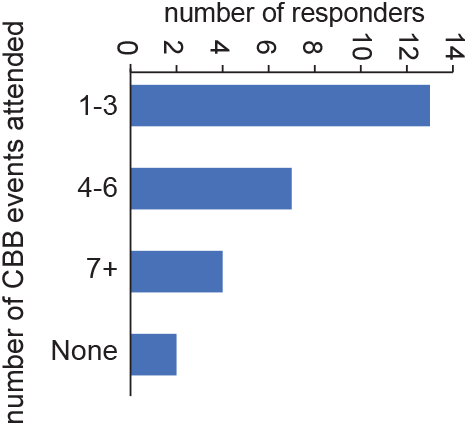
Scripps Community Attendance in CBB-Organized Events. The graph shows the distribution of the number of unique CBB events attended by each survey responder. The data were gathered in 2022 on a sample of the active Scripps community including grad students, postdocs, full-time researchers, staff members, and PIs. The results include all 26 responses from 28 responders who completed this question.

In a follow-up question, we examined the influence of these events on the community’s exposure to bioinformatics. For most of the CBB tools and initiatives analyzed, such as Seminars and Workshops, Discussions, Resources for Troubleshooting, and Networking, the response “Positively” significantly outnumbered “Unchanged,” with an average of 17.25 compared to 10 (**Fig. 6**). Notably, the only CBB initiative that did not have a predominantly positive impact on the community was “Resources for Troubleshooting,” with 11 respondents indicating a positive impact compared to 17 who reported no change.

**Figure 6.**
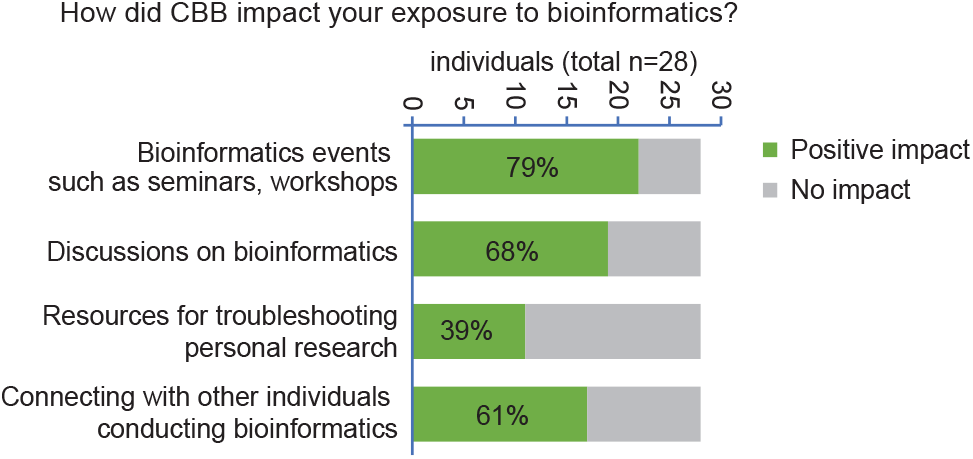
CBB Impact on Exposure Level of the Scripps Community. The graph shows the impact that CBB had on exposing members of the Scripps community to various computational-related activities. The results show a general increase in exposure to 3 out of 4 analyzed topics (Bioinformatics seminars, Discussions on bioinformatics topics, Bioinformatics network), with a total average showing an overall positive impact of CBB. The data were gathered in 2022 on a sample of the active Scripps community (n=28) including grad students, postdocs, full-time researchers, staff members, and PIs. The category “positive impact” includes the following answers: “Significantly more”, “more’, “Significantly more after joining CBB” and “More after joining CBB”; the category “No impact” includes the following responses: “Same exposure”, “Same exposure after joining CBB”.

In our final survey question, we asked participants to rate the usefulness of various CBB features (“Listserv for quarterly updates”, “Listserv for general inquiries and discussion”, “Careers in Bioinformatics Panel”, “Research in Progress Seminars”, “Journal Club”, “Code Topic Discussions”, “One-off Workshops”, “Slack channel for general inquiries and discussions”) across five categories: “Useful,” “Not partaken but interested,” “Indifferent,” “Not useful,” and “Not partaken and not interested.” On average, 13.2 of 28 respondents found CBB features “Useful,” 6.2 expressing interest but being unable to participate, 4.2 indicating indifference, 1.7 finding them “Not useful,” and only 1.1 respondents who had neither participated nor expressed interest (see **Fig. 7**). The seminar series attracted the most interest overall (**Fig. 7**).

**Figure 7.**
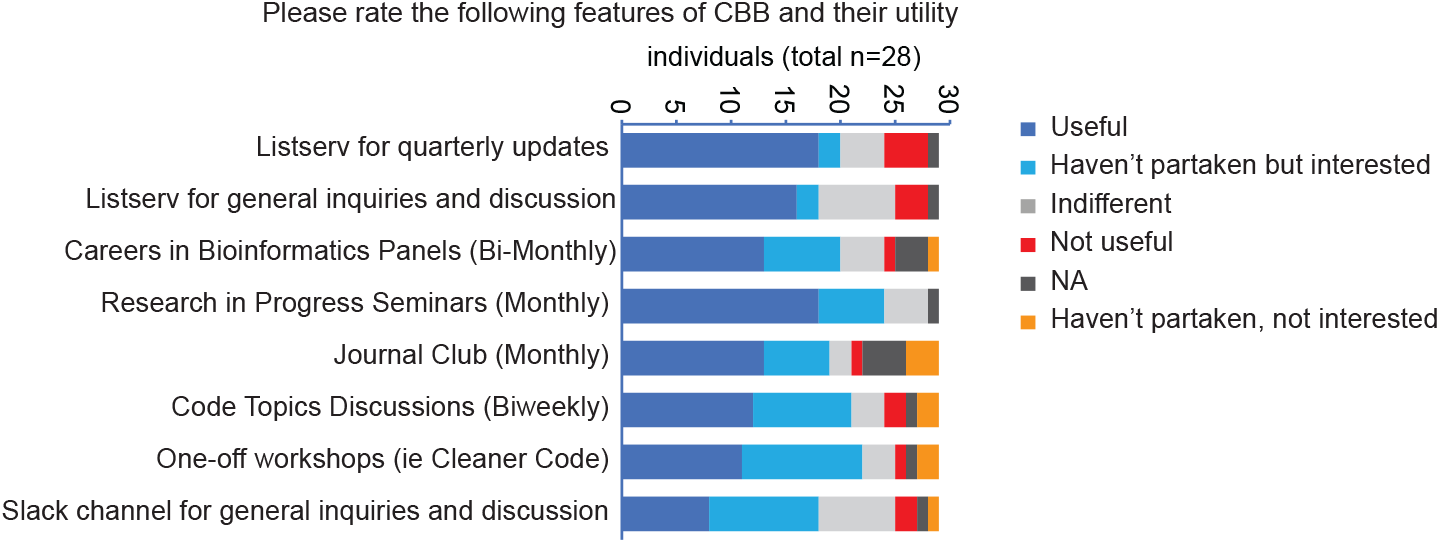
Scripps Community Evaluation on CBB-Organized Events and Tools. The graph shows how the responders evaluated some of the CBB features. NA, not applicable.

Although the sample of 28 respondents is unlikely to be representative of the Scripps community at large, these results demonstrate that CBB activities have had a positive impact on at least some individuals, resulting in increased exposure to information related to computational biology and connecting researchers interested in the field, with most activities deemed useful by those who participated.

These results not only demonstrate the success of the CBB board in engaging the community, with the majority of survey responders finding CBB offerings interesting or useful, but also in organizing initiatives that are well received. The results show a general positive evaluation for all the categories under examination, reflected by the highest average value for “Useful”, followed by “Haven’t partaken but interested” and “Indifferent”. Although we were unable to directly evaluate the effect on computational literacy, these results suggest CBB has increased interdisciplinary community engagement with computational biology and access to learning resources.

## Discussion

Our findings demonstrate that the Scripps Research CBB group has effectively engaged researchers across disciplines, fostering community participation and promoting knowledge exchange in computational biology. Quantitative metrics, including attendance trends and participant feedback, indicate a growing demand for computational training and broad approval of this trainee-led initiative. While direct assessment of the CBB’s impact on bridging specific skill gaps was not feasible—given the open-access nature of events and their diverse scientific focus—the sustained engagement and consistent demand suggest that the group addresses a critical unmet need in postgraduate computational education. Although the survey sample size (n=28) is limited and likely subject to responder bias, the overwhelmingly positive feedback underscores the value of the CBB for a subset of the Scripps Research community, highlighting its role as a complementary mechanism for computational skill development. Importantly, CBB activities addressed the broader need of continuing computational education by increasing exposure to applications of computational biology to real-world research projects and creating opportunities for networking and collaboration.

Analysis of graduate course offerings and core service usage reveals that institutional support for computational research is available through recurring courses and analysis services. However, this support has inherent limitations, particularly for specialized or tailored analyses, due to the constrained variety of computational courses and selective use of custom analysis services. Given the specialized and rapidly evolving nature of computational research methods, institutional resources must prioritize certain approaches over others, limiting their ability to serve as a comprehensive solution.

We do not advocate for a compulsory curriculum like CME for academic researchers but note that the unstructured nature of individual development often leaves low-yield areas unaddressed. Bioinformatic advances can seem irrelevant to non-experts due to perceived low impact or high time costs of implementation but ignoring all technological advances risks loss of innovation and efficiency. The ‘happy medium’ is likely to depend on project details and rate of methodological development in the area. Thus, for promoting workforce-wide bioinformatic competency, approaches beyond compulsory training and complete independent study should be considered^22, 23^. Leaning on communities of practice is one alternative approach that leverages benefits of flexible group/situated learning^24, 25, 26, 27^ and we developed the CBB as a mechanism to augment institutional resources and report evidence supported by data from the CBB case study.

Community-directed programming led by communities of practice offer important advantages over existing institutional resources. Activities led by communities of practice naturally tailor closely to the evolving interests of the group as this is required for their survival^28^. This fluidity overcomes the institutional rigidity of any one department, allowing for exploration of topics that may have traditionally been treated as out of scope.

After trialing multiple event types, we found that the seminar approach was particularly well received, earning the highest approval rating from survey respondents. This likely reflects the relevance of seminar topics to group members, presented in a familiar format that supports varying degrees of engagement. Some seminars provided a deeper dive into tools/technologies and others into the utilization of such tools/technologies to particular fields. Both active and passive participation provide benefits^25^. In contrast, our one-off workshops mostly targeted newer members of the CBB, however this could be expanded in case of sufficient enthusiasm/interest in advanced topics.

An additional benefit of communities of practice transcends the formal programming-related learning opportunities they provide and is tied to the social networks they facilitate. These networks play a large part in setting organizational culture and promoting a sense of belonging^29^. Together these elements are critical to an institute’s identity and help to enhance its constituents’ performance^29^. The survey results show that the CBB’s programming provided opportunities for participants to connect with other individuals (**Fig. 6**), likely supporting these downstream effects.

The limiting factor to community led programming in our experience has been community enthusiasm/interest. The level of this factor is intrinsic to each community. While we show the enthusiasm/interest in the CBB at Scripps Research has been sufficient to support seminar series, we found it was insufficient to support consistent programming for additional series such as journal clubs or workshops. Factors influencing engagement are often beyond the control of organizers, but leadership buy-in is a modifiable factor. Anecdotally, labs that emphasize individual development, both formal and informal, tended to have greater participation. To foster broader engagement, we recommend institutional support of community-driven learning, ensuring that group activities are recognized as productivity-enhancing rather than productivity-competing. Future studies should investigate how communities of practice impact computational skill level and productivity.

## Materials and methods

### Human Subjects

The survey was reviewed by the Institutional Review Board (IRB) of Scripps Health and Scripps Research (IRB-24-8318). The IRB confirmed the survey qualifies for exemption from full review in accordance with article 45 CFR 46.104(d)(2).

### Genomics Core Analysis

We used de-identified data on TSRI Genomics Core experiments over 10 years (2012-2021) exported from an SQL database utilized routinely for experiment tracking. The following fields were used to identify experiments: project_uid, project_name, project_description, primary_investigator_uid, creation_date, sample_type, species. The project_uid was used as a unique identifier of experiments (typically, each discrete experiment would be assigned a unique project_uid). The sample_type column was used to identify experiment types and manually converted into a new sample_type_simplified column to group together experiments of the same type (for example, polyA+ and ribosome depletion RNA-seq library preparation protocols would be grouped under RNA-seq). Experiments not fitting in any major categories were assigned ‘Other_or_NA’ in the sample_type_simplified column. R version 4.2 was used for analysis and plotting.

### High Performance Cluster Analysis

We used de-identified data on the main computing cluster at TSRI over 4 years (2017-2020). Only the following fields were analyzed: Year, Group, CPU days, Number of jobs.

### Seminar Topic Analysis

Seminar information including location, host, attendance, speaker, home lab, seminar title and date delivered were recorded beginning in February 2020. If the presenter was a PI their department affiliation was used and if not, their PI’s departmental affiliation used as the department of speaker. Speakers with unknown departmental affiliations were marked as NA and excluded from the affiliation analysis. Speakers from core services units were assigned an independent department ‘Core Services’. To harmonize diverse department names, categories of research, e.g., neurology => neuroscience, were supplied to “Field Categories”. To designate scientific topic for non-Scripps speakers with unknown Department affiliation their lab website landing page was used to best estimate the broad research category. Only keywords with at least 2 independent seminars were included in the frequency analysis. There were 5 singleton keywords. This yielded a list including: Genomics, Machine Learning, Immunology, Neuroscience, Therapeutics, Structural Biology, High Throughput Computing, Dimensionality Reduction, and Proteomics.

### Survey Data Collection

The survey comprised several questions designed to gauge the extent of community involvement in CBB events and the perceived impact of these initiatives. The primary questions focused on the frequency of attendance at CBB-organized events, the influence of these events on exposure to bioinformatics, and the usefulness of various CBB features.

The survey was distributed on three occasions over a two-month period to mailing lists that included CBB members and early-career scientists (graduate students, postdocs). Utilizing Google Forms, survey data was systematically collected. To uphold respondent anonymity, no personal identifiers were stored. Respondents were explicitly instructed not to submit their responses more than once to prevent duplication.

### Data Analysis

Quantitative analysis of survey responses was performed to discern trends and patterns within the collected data using Python 3, Pandas library. Descriptive statistics were employed to summarize participant demographics, attendance rates at CBB events, and perceptions regarding the impact and utility of CBB initiatives. The frequency distribution of respondents’ attendance at CBB events was analyzed to understand the level of community engagement. The number of events attended by each participant was categorized, and the distribution was visualized using graphical representation. Responses regarding the influence and of CBB events on community exposure to bioinformatics (**Fig. 6**) and rating of CBB features (**Fig. 7**) were analyzed by grouping the responses from a question per each category expressed as bar in the plot. The frequency of positive, neutral, and negative perceptions was tallied for each CBB initiative or feature, and comparative analysis was conducted to identify trends and areas for improvement. Participants’ ratings of various CBB features were analyzed to assess their perceived usefulness. Responses were categorized into “Positively” or “Unchanged” for the services, or distinct ratings, such as “Useful,” “Interested but unable to participate,” “Indifferent,” “Not useful,” and “Not interested,” for the features, and the distribution across these categories was examined.

The freely available version of ChatGPT was used to perform stylistic language editing and proofreading.

## Acknowledgments

The authors would like to thank members of the Computational Biology and Bioinformatics network for their participation in the network activities and survey. We also thank Andrew Su for support and advice, Ian Wilson for departmental support and Pam Graham for administrative support. J. Z. is a recipient of the Cancer Research Institute/Irvington postdoctoral fellowship.

