## SupplementaryInfo for "Catalyzing computational biology research at an academic institute through an interest network"

**Supplementary Table 1. Number of experiments by category performed in the Genomics Core.**

| | Experiment type | 2012 | 2013 | 2014 | 2015 | 2016 | 2017 | 2018 | 2019 | 2020 | 2021 |
|---|---|---|---|---|---|---|---|---|---|---|---|
| 1 | RNA-seq | 52 | 79 | 64 | 135 | 108 | 82 | 111 | 78 | 62 | 60 |
| 2 | DNA-seq | 33 | 22 | 16 | 20 | 16 | 21 | 48 | 88 | 65 | 40 |
| 3 | Other or NA | 28 | 57 | 30 | 28 | 44 | 84 | 89 | 100 | 57 | 45 |
| 4 | ChIP-seq | 10 | 23 | 11 | 16 | 12 | 13 | 7 | 4 | 1 | 4 |
| 5 | miRNA-seq | 5 | 18 | 12 | 11 | 7 | 3 | 4 | 5 | 3 | 1 |
| 6 | Amplicon-seq | 1 | 0 | 5 | 14 | 19 | 22 | 30 | 21 | 12 | 5 |
| 7 | CRISPR | 0 | 0 | 0 | 0 | 0 | 1 | 5 | 22 | 31 | 14 |
| 8 | 10X Chromium RNA-seq | 0 | 0 | 0 | 0 | 0 | 0 | 15 | 36 | 29 | 41 |
| 9 | Lib-seq | 0 | 0 | 0 | 0 | 0 | 0 | 10 | 3 | 44 | 29 |
| 10 | scDNA-seq | 0 | 0 | 0 | 0 | 0 | 0 | 2 | 0 | 0 | 0 |
| 11 | DRUG-seq | 0 | 0 | 0 | 0 | 0 | 0 | 0 | 2 | 18 | 11 |

**Supplementary Table 2. Department affiliations of research groups with largest usage increase vs decrease from 2017-2020.** There is no obvious bias in the department affiliations of labs which increased vs decreased their HPC usage. There is no statistical enrichment of ISCB labs among HPC users who increased vs decreased usage. ISCB, Integrative Structural and Computational Biology; MM, Molecular Medicine; SRTI, Scripps Research Translational Institute; HPC, High Performance Computing cluster.

| | **Usage increase: fold** | **Usage increase: hours** | **Department** |
|---|---|---|---|
| Lab 1 | 4X | 60k | ISCB |
| Lab 2 | 1.2X | 14k | Chemistry |
| Lab 3 | 2X | 12k | ISCB |
| Lab 4 | 6X | 8.7k | MM |
| Lab 5 | 3X | 1.7k | SRTI |
| Lab A | 100% drop | 2.3k | Chemistry |
| Lab B | 100% drop | 7.5k | ISCB |

**Supplementary Table 3. Departmental affiliations of groups which increased and decreased their HPC usage as measured by CPU hours.** ISCB, Integrative Structural and Computational Biology; MM, Molecular Medicine.

| | **Number of labs increased usage** | **Number of labs decreased usage** |
|---|---|---|
| ISCB (excluding emeriti) | 4 | 7 |
| MM | 2 | 2 |

**Supplementary Table 4. Services offered by the CCBB.**

*Scripps Research California core analysis services offered as of 1/11/2024. The most up to date offerings can be found here. ATAC, assay for transposase-accessible chromatin; CCBB, Center for Computational Biology and Bioinformatics; CNV, copy number variant; GEO, Gene Expression Omnibus; NCBI, National Center for Biotechnology Information; SNP, single-nucleotide polymorphism; SRA, Sequence Read Archive; UMI, unique molecular identifier.*

| **Analysis Category** | **Analyses** |
|---|---|
| Transcriptomics | RNA-seq, UMI-based RNA-seq, High Throughput RNA-seq (”DRUG-seq”), smallRNA-Seq (miRNA-seq) |
| Single-cell analytics | Gene expression, Immune profiling (VDJ, VDJ+Gene expression), Epigenome profiling (ATAC, Multiome ATAC + Gene expression), SNP-based demultiplexing of multiplexed data |
| Spatial genomics | Visium Spatial transcriptomics (10x Genomics), GeoMx Digital Spatial Profiling (nanoString) |
| Genomics | Exome, Whole Genome sequencing data analyses, SNPs, indels, CNVs |
| Epigenomics | ChIP-Seq/eCLIP-Seq/CUT&RUN/RIP-Seq, ATAC-seq |
| Metagenomics | 16S rRNA-Seq |
| Custom analyses | Sequencing data analyses for specific lab projects that use standard tools, software packages and/or custom code/script developed by CCBB. |
| Public datasets | Analyses on public dataset available at NCBI’s GEO database repository |
| Functional analyses | Advaita’s iPathwayGuide, Gene Set Enrichment Analysis (GSEA) |
| Consultations | We provide consultations on experimental design, overview of analyses and costs involved. We can take on both short-term (typically less than a month and about 10 hours of CCBB effort) and long-term collaborative projects involving bioinformatic support. |
| GEO/SRA uploads | We can submit sequencing data, related sample metadata and the results of analyses to NCBI repositories like GEO and SRA for a small service fee. |

**Supplementary Table 5**. *Table of all courses offered by Scripps Research (CA and FL) covering computational/bioinformatic topics between 2011-2024.*

| Academic Year | Number of computational/ bioinformatic courses offered | Course titles | Total number of courses offered |
|---|---|---|---|
| 2011-2012 | 3 | Basic Biostatistics, Human Genetics and Genomics, Applied Bioinformatics & Computational Biology | 23 |
| 2012-2013 | 3 | Introduction to Biostatistics, Human Genetics and Genomics, Applied Bioinformatics & Computational Biology | 29 |
| 2013-2014 | 1 | Introduction to Biostatistics | 17 |
| 2014-2015 | 2 | Introduction to Biostatistics, Genetics and Genomics | 29 |
| 2015-2016 | 2 | Introduction to Biostatistics, Applied Bioinformatics and Computational Biology | 22 |
| 2016-2017 | 2 | Introduction to Biostatistics, Genetics & Genomics | 28 |
| 2017-2018 | 3 | Quantitative Data Analysis Bootcamp, Introduction to R, Introduction to Biostatistics | 24 |
| 2018-2019 | 5 | Quantitative Data Analysis Boot Camp, Fundamentals of Scientific Computing, Applied Bioinformatics and Computational Biology, Introduction to Biostatistics, Genetics and Genomics | 33 |
| 2019-2020 | 2 | Quantitative Data Analysis Boot Camp, Introduction to Biostatistics | 24 |
| 2020-2021 | 6 | Quantitative Data Analysis Boot Camp, Fundamentals of Scientific Computing, Applied Bioinformatics and Computational Biology, Advanced Methods in Statistical Analysis, Introduction to Biostatistics, Genetics and Genomics | 30 |
| 2021-2022 | 3 | Quantitative Data Analysis Boot Camp, Introduction to Biostatistics, Computational and Analytical Tools for Chemists | 26 |
| 2022-2023 | 5 | Quantitative Data Analysis Boot Camp, Fundamentals of Scientific Computing, Applied Bioinformatics and Computational Biology, Introduction to Biostatistics, Advanced Methods in Statistical Analysis | 31 |
| 2023-2024 | 5 | Advanced Methods in Statistical Analysis, Introduction to Data Science, Introduction to Biostatistics, Advanced Data Science, Computational and Analytical Tools for Chemists | 32 |

**Supplementary Table 6.** Details on CBB seminar speakers.

| **Career stage** | **Number of speakers** |
|---|---|
| PhD student | 5 |
| Postdoc | 14 |
| Staff scientist | 2 |
| Faculty | 8 |
| Industry scientist | 6 |
| Core facility scientist | 2 |
